# Enhancing volumetric microscopy with blind computational correction of spatially variant aberrations using neural field representation

**DOI:** 10.64898/2026.01.15.699665

**Authors:** Linh Hoang, Zhongqiang Li, Dominique Meyer, Xiankun Lu, Ji Yi

**Affiliations:** Department of Biomedical Engineering, Johns Hopkins University, Baltimore MD, 21231; Department of Ophthalmology, School of Medicine, Johns Hopkins University, Baltimore MD, 21231

## Abstract

Studying biological processes across multi-millimeter scales requires imaging systems that combine high spatial resolution with a large field of view (FOV). However, optical aberrations degrade image quality, particularly in large-FOV systems where distortions gradually worsen toward the periphery. Existing methods for correcting field-dependent aberrations are limited and often impractical. Hardware-based solutions such as adaptive optics demand additional components and sample manipulation, increasing system complexity and experimental burden. Computational approaches, while promising, typically rely on exhaustive point spread function (PSF) calibration or large training datasets, restricting their applicability. We introduce a self-supervised algorithm that simultaneously estimates spatially varying aberrations and reconstructs 3D sample structure from a single blurred image—without PSF calibration or training data. Our method was implemented on optically cleared mouse hippocampal neurons and cortical vasculature imaged with oblique plane microscopy (OPM), as well as in vivo mouse retinal vasculature captured with adaptive optics scanning laser ophthalmoscopy (AOSLO), where PSF calibration is infeasible. The algorithm predicts and corrects aberrations across multi-millimeter FOVs, enhances contrast in low signal-to-noise regions, and reveals critical structural details in neuronal and vascular systems that are obscured in raw images. This approach enables high-resolution, large-scale imaging without hardware modifications, expanding the accessibility of advanced microscopy techniques.

## Introduction

Imaging large volume with high spatial resolution is increasingly called for, because many biological processes or signaling can span over large distance, and rare biological events may be difficult to capture within a small field of view (FOV). Yet, there is an inherent trade-off between resolution and FOV in conventional microscopy systems. Conservation of etendue dictates that the product of the acceptance angle and the imaging stays constant throughout that system. For instance, a high numerical aperture (NA) objective increases resolution by having a larger acceptance angle, but reduces the FOV. Designing sophisticated objectives to support larger etendue yet maintains optical performance would fundamentally address the challenge, however, the complexity and cost can become prohibitive for large scale adoption, and impossible in some applications (e.g. head mounted miniscope, handheld scope or *in vivo* retinal imaging device).

One key factor dictating this trade-off is the optical aberration that deteriorates the resolution when pushing FOV. In addition to sophisticated objectives using multiple elements to minimize aberration, additional hardware elements (*e.g.* deformable mirrors, spatial light modulator) can be introduced to correct aberration via adaptive optics (AO). Direct wavefront sensing AO systems need a point source in the FOV to act as guide-stars, which often are beads injected into the sample^1–3^ or a two-photon point scan^4,5^. Such requirements significantly increase complexity to microscope design and experimental procedures.

In place of the hardware approaches, computational correction methods can break the trade-off to improve resolution regardless of FOV. The most classic method is deconvolution which assumes that aberration is space invariant, which is often untrue especially for large NA and large FOV systems. Patch-wise deconvolution methods^6–10^ deal with field-dependent aberration inherent to most microscopes. These methods divide the image into small patches with apparently invariant aberrations, and deconvolve them with measured local point spread functions (PSF). This leads to a trade-off between accuracy and computation costs with respect to the number of patches. Methods to calibrate the spatially variant PSF as combinations of invariant basis^11–13^ such as Seidel aberrations^14^ reduce the computation costs, but have mainly stayed in 2D images and cannot be incorporated into *in vivo* systems where sample specific aberrations create non-measurable PSFs. Those methods require pre-calibration of the field dependent PSFs - either from the laborious process of characterizing hundreds of thousands of bead measurements, or relying on assumption of rotational symmetry in the imaging system which may not be valid.

Deep learning methods have been developed to reduce computation and eliminate pre-calibration needs. However, networks that rely on training data^15–19^ fail to generalize to other systems. Retraining is prohibitive to users that have not collected the massive amount of data necessary for training. Most networks have been trained on simulated data built on a mathematical model of the image formation process^18,19^, further complicating their ability to extend to real world systems. Self-supervised learning for blind deconvolution circumvents the need for training data and questions of practical relevance^20,21^ but have yet to extend to spatially-variant deblurring. In short, methods to correct spatially varying aberration over large FOV is inadequately addressed.

Here we propose a self-supervised method for field-dependent aberration estimation and correction from a 3D microscope image. We use neural field representation of spatially varying aberrations to impose a smoothness constraint on wavefront variance, and incorporate a physics-informed image formation process in the forward model to exploit the phase information in 3D intensity data. Such approach allows us to predict the field-dependent aberration phase and restore the sample structure without any training data, facilitating application to different imaging systems with as little known as the system NA.

We tested our algorithm on two vastly different microscope systems – oblique plane microscope (OPM) and adaptive optics scanning laser ophthalmoscope (AOSLO) - each with complex pupil function and aberrations. Both modalities pose a challenge to existing deconvolution methods: OPM lacks a clear symmetry in its aberration pattern to be exploited in PSF calibration; AOSLO suffers from sample specific aberrations that are not measurable for calibration; both currently lack a satisfying image formation model with spatially varying aberrations to create simulated data for training. In the OPM, we computationally extend the high-resolution high-SNR region covered by the microscope without costly hardware addition to the system design. In AOSLO, we validated the accuracy of our aberration prediction via comparison with ground truth (GT) wavefront measurements from a Shack-Hartmann sensor. Our method as a post-acquisition processing step significantly improves the capability of these modalities for high resolution, large FOV imaging.

## Results

### 1. Self-supervised algorithm for field-dependent aberration estimation and correction

In any real imaging system, imperfections in optical components such as lenses and mirrors cause aberrations that degrade the image quality. Optical aberrations such as astigmatism and coma vary across the imaging field, usually worsening towards the image periphery (**Fig. 1a**). Aberrations measured at the central FOV rarely describe what happens at the edges, which is why corrections based on a single PSF often fail in large field systems (**Fig. 1e**).

**Fig 1.**
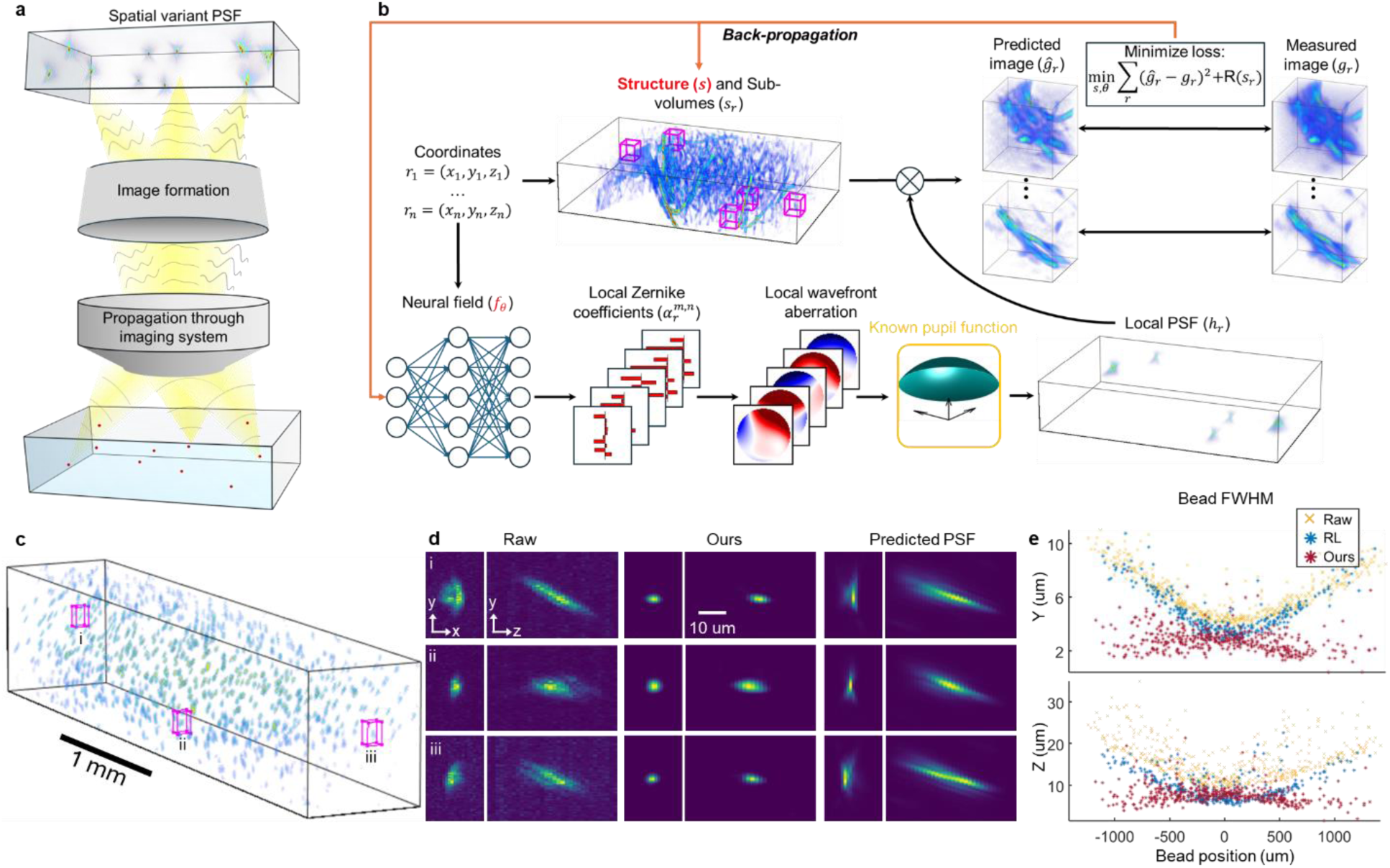
Design concept and characterization. (a) Spatially variant PSF in an imaging system introduced by wavefront aberration through the imaging system. (b) The schematic on our algorithm for predicting the spatially variant aberration and underlying sample structure from a blurry measured 3D image. The forward model requires prior knowledge of pupil aperture function of the imaging system. A neural field representation capture the spatially dependent Zernike coefficients to quantify aberrations. (c) An example of 0.5μm fluorescent microspheres imaged with a custom-built oblique plane microscopy to characterize PSFs. (d) Comparison of PSFs from three measured microspheres marked in c (magenta boxes) between raw, recovered, and predicted PSFs. Two of them are from the periphery of the imaging volume. (e) FWHM of fluorescent microspheres in panel (c) in raw measurement, after Richarson-Lucy deconvolution with theoretical PSF, and after our deconvolution algorithm.

The major challenge to calculating spatially variant aberrations from a blurry measurement is the ill-posedness of the problem - blind Richardson-Lucy deconvolution (RLD) is prone to confusing the PSFs with the sample structure^22^. We use a variety of regularizations and physics-informed priors in our algorithm to limit the solution space of the problem (**Fig. 1b**).

First, we take advantage of the fact that 3D image stacks contain more information about the optical system than 2D images. A 3D intensity image encodes the phase information of an imaging system, which existing phase retrieval algorithms have leveraged to calculate the microscope pupil function from PSF measurements at multiple axial planes^23–25^. Inspired by these approaches, we model each local PSF as the 3D Fourier transform of the microscope’s pupil function, with an aberration phase represented by Zernike polynomials. The pupil function - which defines how much light is transmitted through the system - is usually known for any given microscopic imaging system. Aberration manifests as a wavefront distortion of this pupil function. It is worth noting that the pupil function is in the 3D Fourier space that is reciprocal to the 3D spatial imaging data. Representation of the wavefront as Zernike polynomials constrain the phase to classical optical aberrations and reduce the dimensionality of the solution space compared to pixel-wise parameterization.

Second, we expect the aberrations to vary continuously over the 3D imaging space. To impose this constraint, we model the Zernike polynomials as a neural network (NN) that takes the spatial coordinates (X, Y, Z) and outputs the local Zernike coefficients of said position. The neural field inherently outputs a continuous function over the entire imaging volume^21,26^, enforcing the continuity of the aberration. Since there are fewer optimizable parameters in the neural field than total image pixels, this method also reduces the dimensionality of the problem, allowing faster convergence.

To reduce the memory space, we optimize the structure on a small sub-volume (96×96×96 pixels) each time (**Fig. 1b**). For each sub-volume, the NN predicts a set of Zernike coefficients, which are used to calculate the local aberration phase and subsequently the local aberrated PSF. Convolving the aberrated PSF with recovered structure produced blurred image, which is then compared to the experimental imaging data using a user-defined loss function:

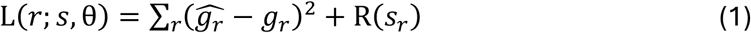

In equation (1), *r* stands for the (X,Y,Z) coordinates of the current sub-volume, *R*(*s*_*r*_) is a regularizer that controls the smoothness and sparsity of the predicted structure *s*_*r*_, *g*_*r*_ is the measured sub-volume and 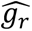 is the predicted blurry image calculated as:

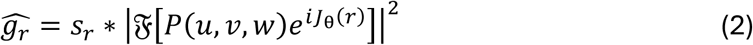

In equation (2), the predicted structure *s*_*r*_ is convolved with the local PSF. *P*(*u*, *v*, *w*) is the circular aperture of the objective lens, *J*_θ_(*r*) is the aberration phase output of the Zernike NN with weights θ, and 𝔉 is the 3D Fourier transform with respect to the pupil coordinates *u*, *v*, *w* in 3D Fourier space. The loss function (1) was backpropagated to update the recovered structure and NN. We randomly computed 20,000 sub-volumes (10 per epoch over 2000 epochs), the sum of which approximately is equal to 100 times the total imaging volume to ensure all 3D space is sufficiently sampled.

**Figure 1c** showed a 3D rendering of 0.5 μm fluorescent microspheres by a custom-built OPM system^27^ – a variant of single objective light sheet microscope - to characterize the 3D PSF. The primary objective is 10x with effective NA of 0.5 (Olympus XLUMPlanFl). The projection views from three representative microspheres across the imaging volume are shown in **Fig. 1d**. Two peripheral microspheres displayed more aberration than the central one. We compared the empirical estimate of the PSF at these three locations to that predicted by the model and observed very similar profiles (**Fig 1d**). Using the aberrated PSF predictions from our network, the algorithm successfully restores peripheral resolution to the same level as center FOV (**Fig 1e**). Using the full width half maximum (FWHM) diameter of the beads as proxy for system resolution, we see that the resolution degrades from around 4.45±0.59 μm lateral and 11.83±2.05 μm axial at center FOV to 9.05±0.74 μm lateral and 22.34±3.77 μm axial at the periphery (**Fig 1e**). The deconvolution using Richardson-Lucy^30^ (RL) using ideal PSF – Fourier transform of the pupil function without aberration phase (**Fig S1a**) – served as a reference to compare the performance of our algorithm (Ours). Non blind RLD with unaberrated PSF does not sufficiently correct aberrations in the periphery; while our algorithm brings both lateral and axial resolution to less than 3.16±1.76 μm lateral and 8.39±3.03 μm axial across the image space (**Fig S1b**).

### 2. Characterizing resolution improvement with optically cleared mouse hippocampus across several millimeter FOV

Large-scale volumetric imaging with high-resolution spatial is essential in to understand connections across different structural scales. We designed an OPM system^27^ that is capable of a mesoscopic FOV (e.g. 3.2 mm x 3.2 mm) with sub-cellular resolution in 3D. Pushing FOV does comes with a price that the aberration became apparent towards image periphery. We set out to test the validity of our algorithm in recover the resolution and predict the aberration across the imaging volume.

In this system, an oblique light sheet illuminates the sample, and a remote focusing objective lens (OL3) placed at an angle from the primary objective (OL1) refocuses the slanted image plane onto the camera (**Fig 2a**). Because OPM uses two objectives whose pupils overlap at an angle, we extend our model to predict aberrations for each pupil separately (**Methods 1a**). This flexibility of incorporating custom imaging models allows the same framework to accommodate more complex imaging geometries than standard epifluorescence microscopes.

**Fig 2.**
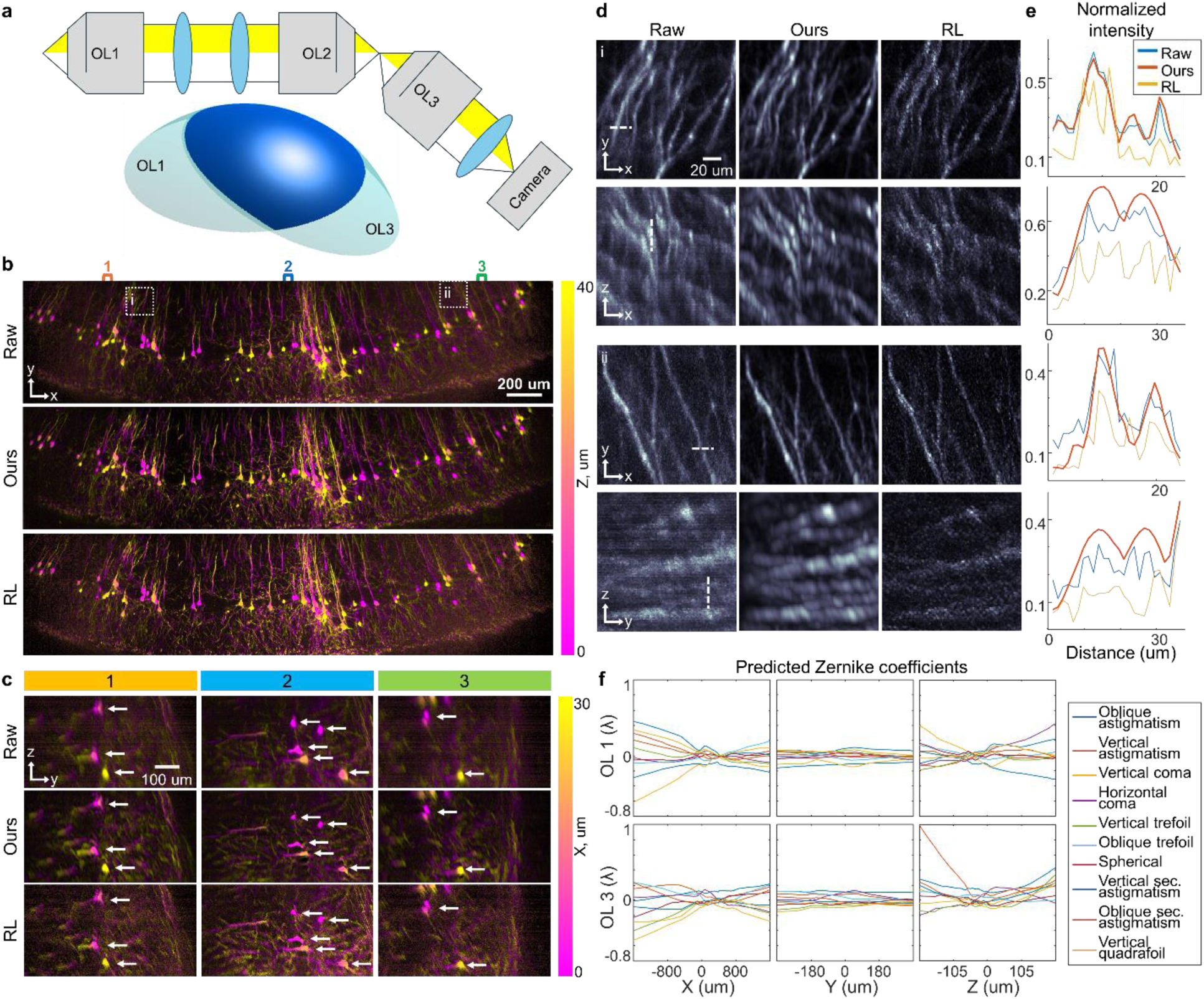
3D imaging of optically cleared mouse brain captured with oblique plane microscopy. (a) A simplified OPM system layout and 3D pupil function. OL=objective lens. (b) Color-encoded *en face* view of the raw and deconvolved image on C57BL/6. Richarson-Lucy (RL) deconvolution is used as comparison with our algorithm. (c) Color-encoded side view of three different image positions marked with colored brackets in panel (b). (d) Maximum intensity projection of two small ROIs in panel (b) (white squares) showing neuronal processes. (e) Pixel intensity across two or more dendrites indicated by white dashed lines in panel (d). (f) Predicted optical aberrations across the image space.

**Figure 2b** showed a color-encoded *en face* projection of imaging volume from mouse mouse hippocampus transgenically labeled Thy1-EGFP, which is often used to track the number, length, and shape of neuronal projections^28^. For applications in neural circuit tracing, the ability to discern individual axons and dendrites over a large area is important in mapping the neural pathways underlying brain functions^29^, making uniform resolution across the FOV essential. The entire 0.5 gigapixels (GP) imaging volume is 3.2mm x 0.7mm x 0.3 mm. A 40μm layer in Z is presented for better visualization.

We took three examples of side projection (y-z projection) from central field and two peripheral fields (**Fig. 2c**). The hippocampus neurons show the apical dendrites as a single long, thick dendrite with several branches leading from the cortical surface to the pyramidal soma, from which axons and basal dendrites with profuse branching arise. The effect of deconvolution is most aggressive in the axial direction as seen from the reduced neuron soma in **Fig 2c** (white arrows). This is consistent with the shape of the measured PSF, which is more elongated axially than laterally (**Fig 1d**). While individual somas are identifiable even in the severely aberrated image periphery, dim and dense dendritic processes become difficult to trace without correction. Our method restores these structures across the entire FOV.

To illustrate the improved visibility of dendritic branching in the image deconvolved with our algorithm, we examine two areas of apical dendrites at the image periphery (**Fig 2d**). The increased resolution reveals arborizations (**Fig S2**, red arrows) that are not noticeable in the raw and RLD images, as well as facilitate detection of adjacent parallel dendrites (**Fig S2a**, white arrows). Such improvements assist in efforts to map neural connections from large scale image data. The deconvolved FWHM of dendrite cross-sections in **Fig 2e** approach the deconvolved system resolution (3.91±1.20 μm lateral, 8.95±0.74 μm axial).

We examined the predicted Zernike coefficient by NN over the imaging volume (**Fig. 2f**). We plotted 10 aberrations from OL1 and OL2 changing in three dimensions. As characteristic of NN outputs, the Zernike NN predicts aberration coefficients that vary continuously across the FOV. Notably, we were able to observe and confirm astigmatism – the most prominent predicted aberration along Z axis in the refocusing objective – in our OPM system by changing the image plane position along Z (**Fig S3**). We also observed the aberration varies more within the light sheet plane in x, z dimension, than the y. OPM collects one 2D oblique oblique light sheet frame at a time and scans the light sheet through the volume to compose 3D image. Signals within oblique frame pass through different parts of all optics, while the de-scanned signals along the scanning Y direction pass through the same areas in the remaining components. Consistent with the physics of optical systems, predicted aberration remains consistent along the galvo-scanning axis Y, and increases further away from image center on the widefield axis X and Z (**Fig 2f**).

When deconvolving different imaging data taken from the same system, the Zernike NN trained on one dataset can be reused as initialization on a different data to speed up computation. For example, we reused the Zernike NN weights trained on this mouse brain image to deconvolve a microsphere image taken with the same OPM system (**Fig 1c**) and achieved excellent results (**Fig 1e**). Although the Zernike NN was trained only on the biological data and not the bead data, the field-dependent PSFs predicted from the mouse brain image closely resembled the spatially varying microsphere patterns imaged with the same optics (**Fig 1d**), validating the accuracy of our aberration prediction.

### 3. Validating structure prediction and characterizing contrast improvement on mouse brain vasculature imaged on OPM

As seen in **Fig 2**, our deconvolution algorithm reveals peripheral features that are not readily detectable in the raw data. We reasoned that the algorithm reassigns scattered signal back to its correct spatial origin, effectively boosting signal-to-noise ratio (SNR) in regions where aberrations are strongest. To further validate, we confirmed the accuracy of these reconstructed features on a different optically cleared brain sample with a ground truth where the peripheral structures were imaged with low aberrations. We also used a higher magnification objective to test the validity of the algorithm with different optics.

We switched the objective in the OPM system a 20X immersion objective (Zeiss Clr Plan-Neofluar) with NA=1.0, which we used to image the frontal cortex of a whole mouse brain optically cleared to a refractive index of 1.49, expressing fluorescence in the endothelial cells lining the blood vessels, which make up the cerebrovasculature. Investigation into the integrity of the cerebral vascular structure is common in studying, diagnosing, and monitoring of brain pathologies^31–33^. **Figure 3a** shows a color-coded *en face* projection of the 2GP 1.6mm x 1.0mm x 0.1mm imaging volume before and after deconvolution. Major arterial branches heading along the brain midline are visible in the raw image, but many finer vessels supplying the frontal lobe – especially near the edges – are obscured by aberrations and vignetting (due to limited lens diameter or sample curvature).

**Fig 3.**
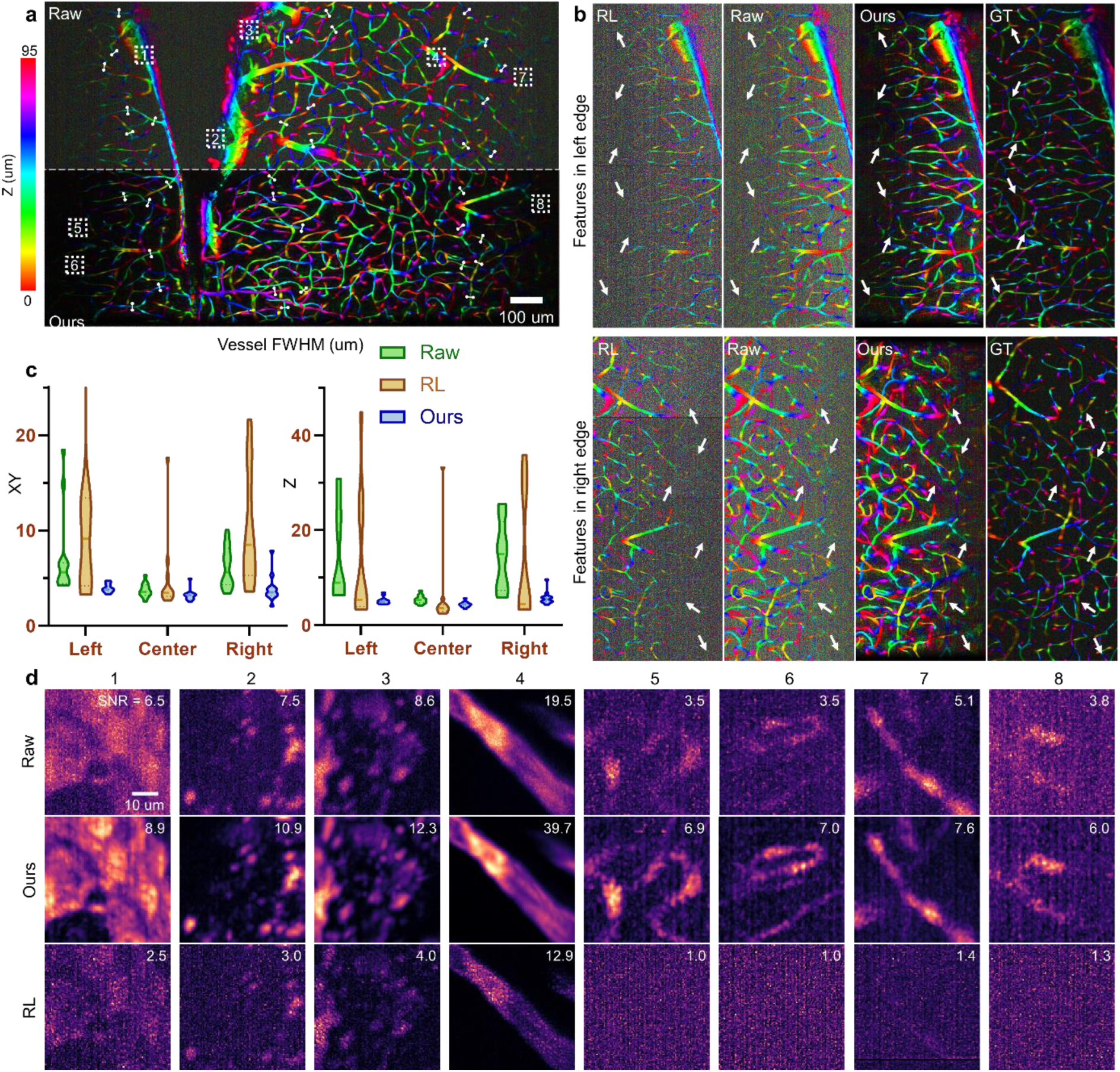
3D image of the mouse brain vasculature captured with an OPM. (a) Depth-projected enface view of the raw and deconvolved brain image. (b) Contrast-adjusted display of the left and right edges of the image in (a) to show the vignetted features brought out by deconvolution (white arrows), validated with high-resolution GT. (c) Axial and lateral diameters of selected vessels in a (white lines) at image center and peripheries (d) Maximum intensity projection of 8 small ROIs in a (white squares). The raw images underwent 2×2×2 binning to mimic the effect of TV regularization in RLD and our deconvolution methods.

The forward model for the 20X objective differs from the 10X model in the size of the circular aperture *P*(*u*, *v*, *w*), since the two objectives have different NA, but the overall framework remains unchanged. Our method reveals vessels in image peripheries that are invisible in both the raw and RLD images (**Fig 3b**). To confirm that these features are not artifacts from the deconvolution algorithm, we obtain two GT images of the peripheries by mechanically moving the sample stage laterally to bring the edges of the original image into the low-aberration center FOV. The vessels revealed by our algorithm matched those in the GT images (**Fig 3b**, white arrows), validating the accuracy of the recovered structure in our algorithm.

We attribute the improved contrast to the ability of our algorithm to consolidate aberrated signals that would otherwise appear as noise into a single originating pixel. Standard RLD sharpens vessels near the center but worsens image quality in the highly-aberrated periphery, where applying the ideal PSF is inappropriate (**Fig 3c**). In contrast, our algorithm restores the peripheral resolution to near the same level as center FOV.

To quantify the improvement in image contrast offered by our algorithm, we compare the SNR across multiple regions of interest (ROIs) spanning the image center and periphery (**Fig 3d**). To isolate effect of signal recovery by deconvolution from the smoothing effect of TV regularization in the loss function, we applied 2×2×2 binning to the raw data before SNR calculation. Our algorithm increased SNR by 40% to 100% relative to the binned raw image, as opposed to RLD which decreases SNR by amplifying noise (which is a well-documented effect of this class of division-based deconvolution^34^).

### 4. Wavefront prediction with AOSLO imaging of in-vivo mouse retina

Retinal imaging is a valuable tool for the study, diagnosis, and management of retinal diseases and vision science. However, imaging the retina is fundamentally limited by optical aberrations introduced by the cornea and lens, which vary from eye to eye and distort the image. Such aberrations cannot be pre-calibrated with a PSF measurement, as there are no non-destructive methods for placing sub-resolution targets like beads inside the eye. The state-of-the-art solution is the addition of a wavefront sensor and a deformable mirror (DM) to measure and compensate for aberrations, which is called adaptive optics (AO). However, the DM can only compensate within an isolanatic patch over an area of less than 2.5° where aberrations is considered are constant^35,36^. As a result, AO improves only a small fraction of the FOV, leaving most of the image (30°-50° viewing angle on a regular fundus camera) uncorrected. Repeatedly measuring and correcting local aberrations at each image location^37,38^ is possible, but it would slow the imaging process and exacerbate motion artifacts. Using multiple DMs or phase modulators^39–41^ can expand the corrected area, but adds to the cost and complexity of the system. Such application is where the ability of our algorithm to predict and correct for spatially variant aberrations in post-processing would be valuable.

To compare our algorithm performance to AO correction, we imaged *in vivo* mouse retina with 0.1ml 2.5% FITC-dextran solution retro-orbitally injected to visualize the retinal vasculature with a previously described AOSLO system^42^. SLO is analogous to confocal microscopy that uses ocular optics instead of microscopy objectives. Sample-specific refractive errors such as astigmatism reduce both resolution and signal strength of the SLO system. With AO, a Shack-Hartman wavefront sensor (SHWFS) measures aberrations and applies a compensatory wavefront to a DM for correction. Depth-scanning for 3D imaging is achieved, in our case, by applying different defocus phases to the DM. In wide FOV systems such as ours (viewing angle ∼20°) the spatial variance of ocular aberrations poses an additional challenge in wavefront correction, since the SHWFS only measures the wavefront within a 4° angle patch at a time. An AO-corrected image only showed high SNR and resolution near the region where wavefront sensing was performed (**Fig S4**).

**Figure 4a** shows an *en face* maximum intensity projection of the non-AO SLO 3D image, presenting large arteries and veins branching out from the optic disc and smaller vessels in the capillary. In comparison, **Fig. 4b** shows the predicted structure from non-AO image by our deconvolution algorithm. The entire 0.26 GP imaging volume is 1.1mm x 1.1mm x 0.2 mm. In provide ground-truth, we applied SHWFS at five retinal locations to measure ocular aberrations. Applying each of these measured wavefronts to the DM, we also collected five locally AO-corrected 3D images, one for each wavefront sensing area (**Fig S4**, wavefront areas marked by asterisks). In each AO-corrected image, capillaries appeared brighter and more defined within the AO location. For each AO dataset, we selected the region with the clearest capillary structure and identified the corresponding regions in the non-AO and reconstructed images.

**Fig 4.**
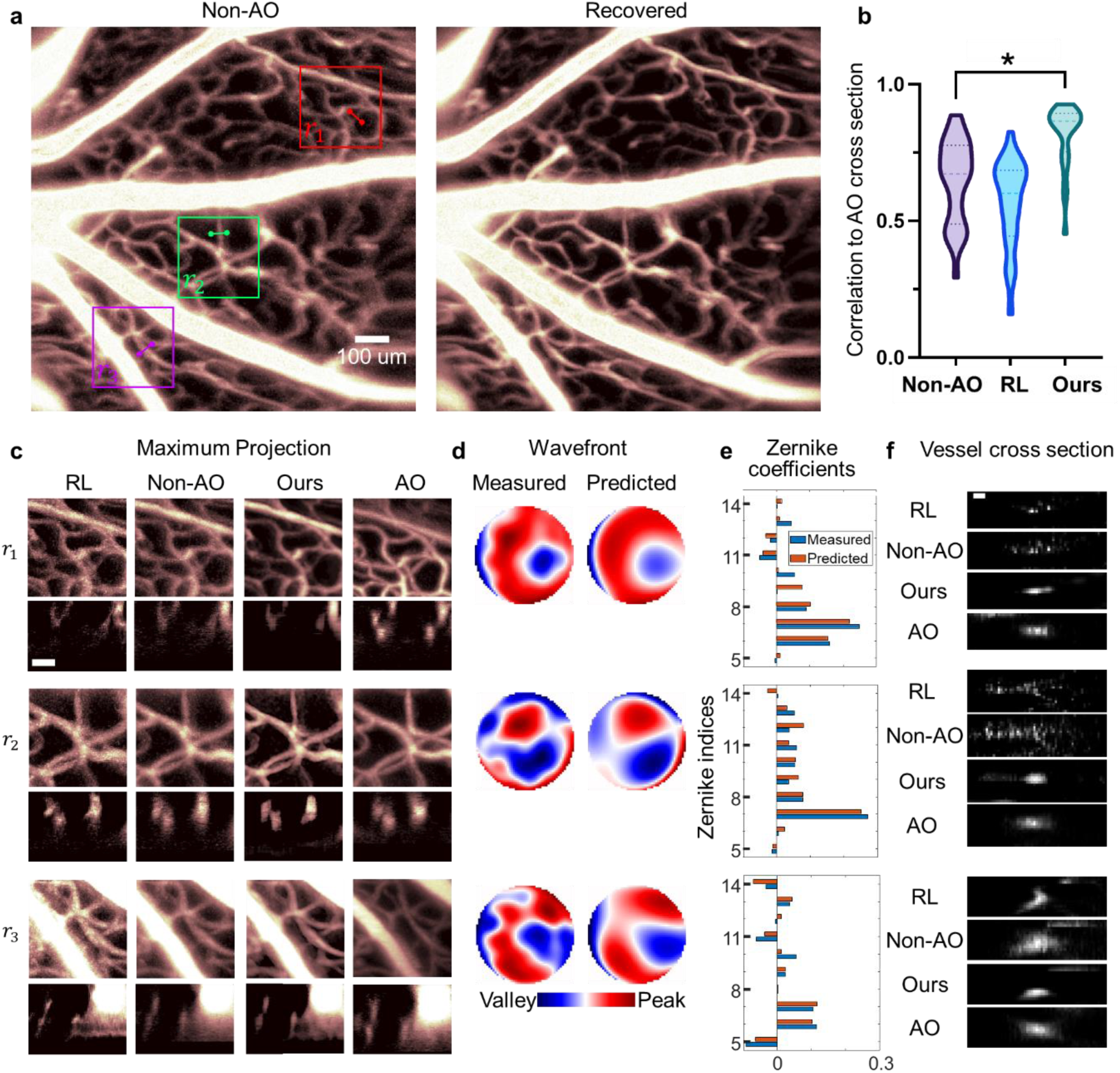
Image of in-vivo mouse retina captured with AOSLO. (a) En face maximum intensity projection of 3D non-AO SLO image and predicted 3D image by our algorithm. (b) Correlation between the cross section of selected vessels in non-AO, RLD, our deconvolved images with AOSLO measurements. *p<1e-7 (c) *En face* and side view of select ROIs in a (colored squares) from non-AO, RLD, our deconvolved image, and AO-corrected measurements. Scale bar: 50 um. The experimental SHWFS measurements and aberration predictions for these ROIs are shown as the phase map (d) and Zernike coefficients (e). (f) Cross section of selected vessels in panel a (colored lines). Scale bar: 1 um.

With the ground truth, we compared the cross section of 25 select capillaries in the non-AO and deconvolved images with locally AO-corrected images (**Fig S5**). Our reconstructions showed the highest correlation with AO-corrected images (0.82 ± 0.12), outperforming both the non-AO data (0.64 ± 0.15) and RL deconvolution (0.57 ± 0.16) (**Fig 4b**). Corresponding ROIs for the wavefront measurements are presented side by side for comparison in **Fig 4c** and **S4b**. Across all ROIs, our predicted structure shows improved resolution and contrast comparable to AO correction – without requiring multiple wavefront measurements or repeated image acquisition.

We then compared our predicted aberration wavefronts with the experimental SHWFS measurements (**Fig 4d, S4c**) from the ROIs in **Fig 4c** and **S4b**. Even though our model predicts only the first 10 Zernike modes (vs. 33 measured by SHWFS), the predicted wavefronts closely matched the measured ones, with an RMSE of 8.56% ± 1.99% of the wavefront amplitude. The predicted Zernike coefficients also closely matched SHWFS values, correctly identifying primary astigmatism and coma as the dominant aberrations (**Fig 4e, S4d**).

**Fig 4f** shows the cross sections of a select blood vessel in each of the 3 ROIs in **Fig 4c**. In *r*_1_ and *r*_2_, the capillary signals are severely scattered by aberrations that RLD with theoretical PSF completely failed to recover the vessel shapes, while our deconvolution effectively collected the scattered signals that closely resembles that in the AO-corrected cross section. In *r*_3_ where aberration is less severe and the vessel only appears enlarged, RLD still failed by over-deconvolving into a structure uncharacteristic of blood vessel morphology, while our algorithm once again matches AO correction. The pixel-wise correlation of these cross-sections (along with 22 other vessels plotted in **Fig S5**) are the statistics plotted in **Fig 4b**.

## Discussion

Using a neural representation of the field-dependent aberrations, we developed an algorithm that simultaneously predicts the spatially variant PSF of an imaging system and the underlying 3D structure of the sample. By incorporating the physics of image formation into the forward model, our method forgoes the need for a calibration image in PSF prediction, making it suitable for *in vivo* systems where significant aberrations come from the sample, and collection of calibration image is impractical. Our self-supervised method does not require external training data, making it readily adaptable to any imaging modalities with as little information as the system NA.

Extensive algorithms have been developed for spatially variant and blind deconvolution, though rarely together. Existing approaches tend to require PSF measurements^6–14^, synthetic data from microscope simulators^18,19^, or real data from multi-modal imaging systems with both high- and low-resolution outputs^15–17^. Otherwise, they rely on assumptions regarding spatial invariance^20,21^, or symmetry in the imaging system^14^, which may not apply to more complex modalities such as OPM or sample-based aberrations such as the cornea. To our knowledge, our algorithm is the first to allow for both calibration-free aberration correction and field-dependent PSF prediction without training data or restrictive assumptions.

We demonstrated the capability of our algorithm to recover aberration wavefront and sample structure on diverse imaging systems and samples with complex PSF (OPM) and aberrations (AOSLO). We validated the predicted aberration using two different methods: by comparison with measured PSF from sub-resolution beads in OPM, and by comparison with Zernike coefficients and aberration phase from SHWFS. Similarly, we validated the recovered structure by comparison with images from low-aberration areas in the same system, and with AO-corrected images.

Our algorithm predicts and corrects aberrations across widefield FOV and improves contrast in low SNR areas due to vignetting or weak fluorescence signals. In OPM, near three-fold improvement in resolution and two-fold improvement in SNR were achieved in the image periphery. In AOSLO, the aberration-corrected viewing angle was extended four-fold. This would be particularly useful for fast volumetric imaging of large samples, since we only need a single image for aberration estimation, as opposed to digital AO which requires multi-view measurements^43,44^. This algorithm also has the potential to increase penetration depth by correcting depth-dependent aberrations in thick samples. Our method would increase the throughput of imaging techniques, thereby advancing quantitative biological studies of large scale dynamic systems, such as neuronal activities or vascular kinetics.

Further characterization of the performance limits of our algorithm is desirable, such as the minimum SNR or maximum aberration strength required for successful reconstruction. We are also investigating its performance on non-sparse samples, such as in transmission or reflectance microscopy. For future development, applying neural representation to the sample structure may reduce the memory footprint of our algorithm, allowing for aberration correction of larger dataset. Another avenue for improvement is parallelizing the deconvolution process over multiple graphical processing units (GPU) to reduce run-time, facilitating the application of our method on time-series volumetric data. If possible, a more mathematically rigorous formula for convolution with spatially variant PSF than our current method of piece-wise convolution could be used for the forward model of image formation to improve reconstruction accuracy.

## Methods

### 1. Joint aberration estimation and structure recovery

Code is designed and developed with PyTorch and is publicly available at.

#### a. Zernike NN for field-dependent aberrated PSF

We employ a multi-layer perceptron (MLP) to represent the Zernike aberration coefficients as a function of image coordinates. The MLP *J*_*θ*_ is defined by a set of parameters denoted as θ, representing the weights of the neural network. The MLP takes a set of 3D image coordinates r = (*x*, *y*, *z*) and outputs 10 Zernike coefficients *a*_*n*_ at Noll indices *n* for each coordinate. Based on previous works where using more than 10 Zernike modes reduces performance^21^, we use Zernike modes 5-14, excluding the first 4 modes representing piston, tip-tilt, and defocus (**Table S1**):

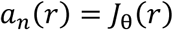

The cumulative aberration at **r** is then:

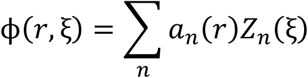

In equation 4, *Z*_*n*_(*ξ*) is the Zernike mode *n* at pupil coordinates ξ = (*u*, *v*). The PSF of fluorescence microscopy at **r** is the 3D Fourier transform 𝔉 of the imaging system aperture *P*(*ξ*, *w*) with respect to the pupil coordinates (*u*, *v*, *w*) with the aberration phase *ϕ* applied:

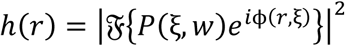

In the case of OPM, the MLP outputs 20 Zernike coefficients for each input coordinate, 10 for OL1 pupil function and 10 for OL3 pupil function (**Fig 2f**). To simulate the effect of an oblique shift in *y* axis at the remote focusing plane, OL3 normalized pupil coordinates are shifted by 1 unit in the *v* axis compared to OL1 pupil. The cumulative aberration of an OPM system is:

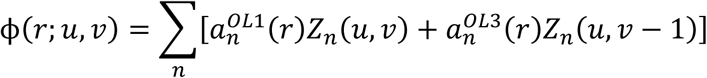

The aberration mask is first calculated with a generously padded array to avoid Fourier transform artifacts in the resulting PSF. The PSF array is then down-sampled by sum-pooling to match the pixel size of the input image.

#### b. Image formation forward model

From the full predicted structure *s*, we crop a small sub-volume *s*_*r*_ centered at **r**. During gradient update, the predicted structure will end up with negative pixels. To impose non-negativity constraint, we apply GELU activation function to *s*_*r*_, which brings negative pixels to zero without cutting off the gradient flow to these pixels as does RELU. The reconstructed blurry image is a convolution of *s*_*r*_ with its corresponding PSF, which we replace with a Fourier space multiplication to reduce computational cost:

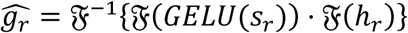

The size of each sub-region is empirically chosen to be at least double the size of h, so that we can discard the edge of 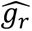 to avoid wrapping artifacts. Each epoch 10 random sub-regions are used, and the total number of epochs is chosen such that on average each pixel of the image is visited 100 times.

#### c. Loss function

The loss function compares the reconstructed blurry sub-region 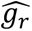 with the corresponding sub-region *g*_*r*_ from the experimentally acquired input image (Eq 1). We updated both the underlying structure *s*_*r*_ and the Zernike NN weights θ to minimize the mean squared error (MSE) between prediction and measurement. Aside from the fidelity term, a regularizer ensures the structure converges to an expected fluorescence image with sparse and smooth signals:

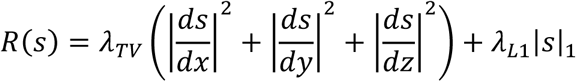

When the regularization rates *λ*_*TV*_ and *λ*_*L*1_ are constant, the loss function tends to be dominated by the MSE term at early epochs and then by the regularizer at later epochs. This caused the Zernike NN to quickly converge to a nonoptimal local minimum, and the predicted structure to be oversmoothed when the regularization dominates. To maintain balanced proportions of fidelity and regularization in the loss function throughout the learning process, we detach each term from the gradient tree and calculate a regularization rate that scales the TV and L1 terms to a fixed fraction of the MSE term. For example, if in one epoch the fidelity loss is 1e-4, the TV loss is 1e-3, and the regularization is set to be 0.1 of the fidelity, then *λ*_*TV*_ = 0.01 for this epoch. The rates *λ*_*TV*_ and *λ*_*L*1_ are updated every epoch.

Optimization is performed using the RAdam optimizer with β1 = 0.9, and β2 = 0.999. The learning rates (LR), number of epochs, and regularization parameters for each image in the paper are reported in **Table S2**. The algorithm ran on a NVIDIA T4 GPU.

### 2. Experimental details

All animal-related procedures including mouse whole-brain clearing and mouse in vivo imaging were in accordance with the Institutional Animal Care and Use Committee at Johns Hopkins University and conformed to the guidelines on the Use of Animals from the National Institutes of Health (NIH). Animals were maintained on a C57BL/6 background under approved protocols.

#### a. Cleared mouse brain imaging

We used a previously published OPM setup and clearing protocol^27^.

A specimen expressing Mobp-EGFP was stained for the Thy1 CreERT2 cell line (**Fig 2**). For clearing, brains were perfusion-fixed in 4% PFA, washed in PBS at 37 °C, delipidated in CUBIC-L with periodic solution refreshes, and refractive-index-matched in CUBIC-R+ until samples were optically homogeneous. Samples were then secured in 30 mm dishes with UV-curable adhesive and immersed in the same matching medium. For vasculature visualization, we used Tie2-Cre;Ai9 double-transgenic animals (endothelial Tek/Tie2 promoter–driven Cre crossed to tdTomato reporter), which yield tdTomato labeling of endothelial cells (**Fig 3**). Tissue was stabilized with SHIELD OFF/ON chemistry, delipidated in CUBIC-L, and finally index matched in a modified high RI uRIMS formulation tuned to RI ≈ 1.496 at pH ≈ 7.4. The bath was sealed with sunflower oil to prevent evaporation induced crystallization of the iodixanol solution.

Two different primary objectives were used for OPM imaging. The neuron-stained hippocampus was imaged under a 10X NA=0.6 water immersion lens (Olympus XLUMPlanFl). The endothelium-stained cortex was imaged under a 20x NA=1.0 clearing medium immersion lens (Zeiss Clr Plan-Neofluar). Operating wavelength was 488nm for illumination and 509nm for detection. Acquisition control was managed via custom LabVIEW software. Y-axis scanning for volumetric imaging was performed either with a galvo mirror (Nutfield QS-12 OPD) for neuronal brain or a motorized stage (Thorlabs Z825B) for vascular brain.

#### b. Mouse retina imaging

We used a previously published AOSLO setup and animal handling protocol^42^.

Throughout the imaging process, a mixture of 1% (v/v) isoflurane and oxygen was administered to maintain stable anesthesia, while a heat pad kept the body temperature at 37°C. After the mice were dilated and anesthetized with ketamine/xylazine, 0.1 ml of 2.5% Fluorescein isothiocyanate–dextran (46944-500MG-F, Sigma-Aldrich Co) was injected retro-orbitally to enable visualization of the retinal vasculature. A +10 D contact lens with hypromellose gel was applied to the cornea to prevent dehydration and cataract formation during imaging. The mice were then positioned on the imaging stage for in-vivo retinal imaging using the DualCH-AOSLO (**Fig 4**).

The AOSLO system uses a series of galvo mirrors for 2D scanning across the retina, and a DM (ALPAO, DM97-15) for depth-scanning as well as wavefront correction. A multimode fiber serves as the detection pinhole. A quarter wave plate and polarizing beam splitter separate the light returning from the retina from surface reflections from relay lenses. For AO imaging of fluorescein or GFP, a 488 nm laser was activated for wavefront sensing, focusing on imaging planes around the inner vascular plexus (IVP) layers by adjusting a tunable lens (ML-20-37, Optotune Switzerland AG). The AO closed loop was then initiated to correct for aberrations. The custom acquisition software was developed in C++, integrating MATLAB Engine to control the SHWFS and DM.

### 3. Image processing

#### a. Bead characterization

We characterized the 3D resolution pre- and post-deconvolution for OPM with 10x objective using 0.2 µm fluorescent microspheres embedded in solidified agarose gel (**Fig 1c-e**). Bead image was denoised with a gaussian filter before segmentation with adaptive Otsu thresholding. Bead centers were detected as the local maxima in foreground regions. Beads within a 25-pixel radius of the image edges or another bead were removed. A volume of size 20×50×40 pixels cropped around the bead center defines the experimentally measured PSF. Volumes with brightest voxel over 15 pixels away from the center were removed. The remaining beads were linearly interpolated in 3D to increase the pixel density 4 times. A gaussian curve is fitted to the 1D maximum projection of the bead in 3 axes, and the FWHM of the PSF is calculated from the gaussian fit parameters. Beads where gaussian fitting failed or returned a FWHM larger than the PSF array size due to noise were removed. A total of 730 beads were characterized (**Fig 1e**). The beads were ordered with respect to their position in x axis, and a sliding window of 100 beads was applied to calculate the moving average and standard deviation (STD) of the bead FWHM across the lateral FOV (**Fig S1**).

#### b. Vessel characterization

For 20x OPM and AOSLO images, user-selected vessels were manually traced in en face view with a straight line from one branching point to another, and the cross section of each vessel is the pixel intensity along a line perpendicular to the traced vessel at its midpoint (**Fig 4a**). The cross-section line does not necessarily follow the image axis, so 2D linear interpolation was used to obtain the pixel intensity of said line in each z frame (**Fig 4f**). For 20x OPM, FWHM of the vessel is calculated from gaussian fitting to the 1D maximum projections of the cross-section in x-y or z (**Fig 3c**). For AOSLO, corresponding vessels were manually identified in images with local AO correction, and vessel cross-sections in AO images were obtained as described above. Cross-sections in AO and non-AO images were registered using cross-correlation, and pixel-wise Pearson correlation coefficients of gray values between cross-sections of corresponding vessels were calculated (**Fig 4f**).

#### c. SNR characterization

Selected ROIs in the OPM vascular brain image were segmented by manually chosen thresholding. Morphological opening and closing, and removal of isolated pixels were applied to clean up the segmentation. SNR is the ratio of the average signal pixel intensities over the STD of background pixel values (**Fig 3d**).

## Acknowledgement

This work was supported by fundings from NIH, EB034272, EY032163, EY034607, Boons Pickens endowment fund, Wilmer unrestricted grant from Research to Prevent Blindness. Dominique Meyer is supported by Kavli doctoral fellowship.

