## Supplemental Materials for "Enhancing volumetric microscopy with blind computational correction of spatially variant aberrations using neural field representation"

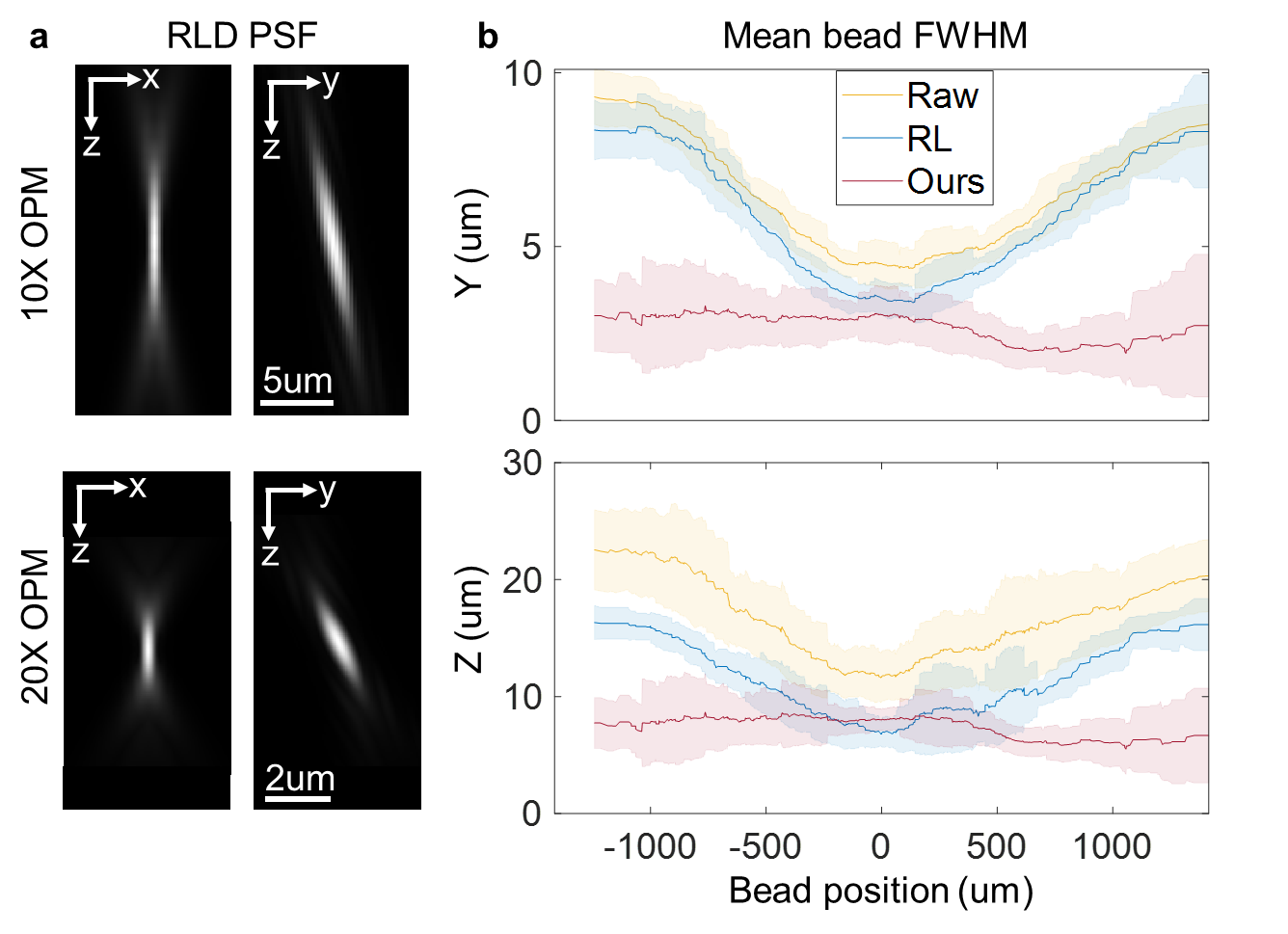

**Supplemental Figure S1.** (a) Unaberrated PSF used for RLD. (b) Moving average and STD of bead lateral (y) and axial (z) FWHM against image position (x).

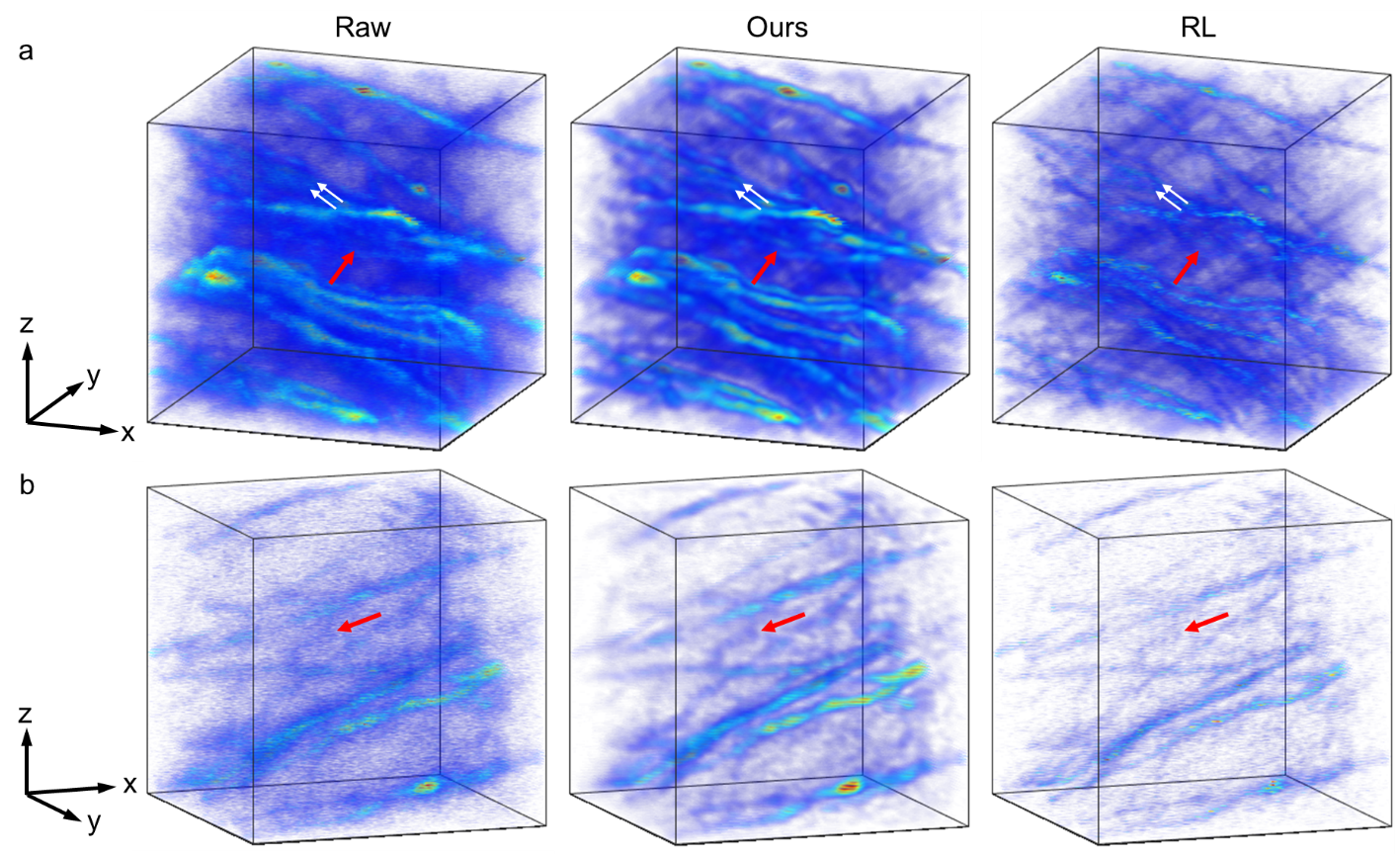

**Supplemental Figure S2. 3D rendering of mouse hippocampus neuron ROIs shown in (a) Fig 2di and (b) Fig 2dii.**

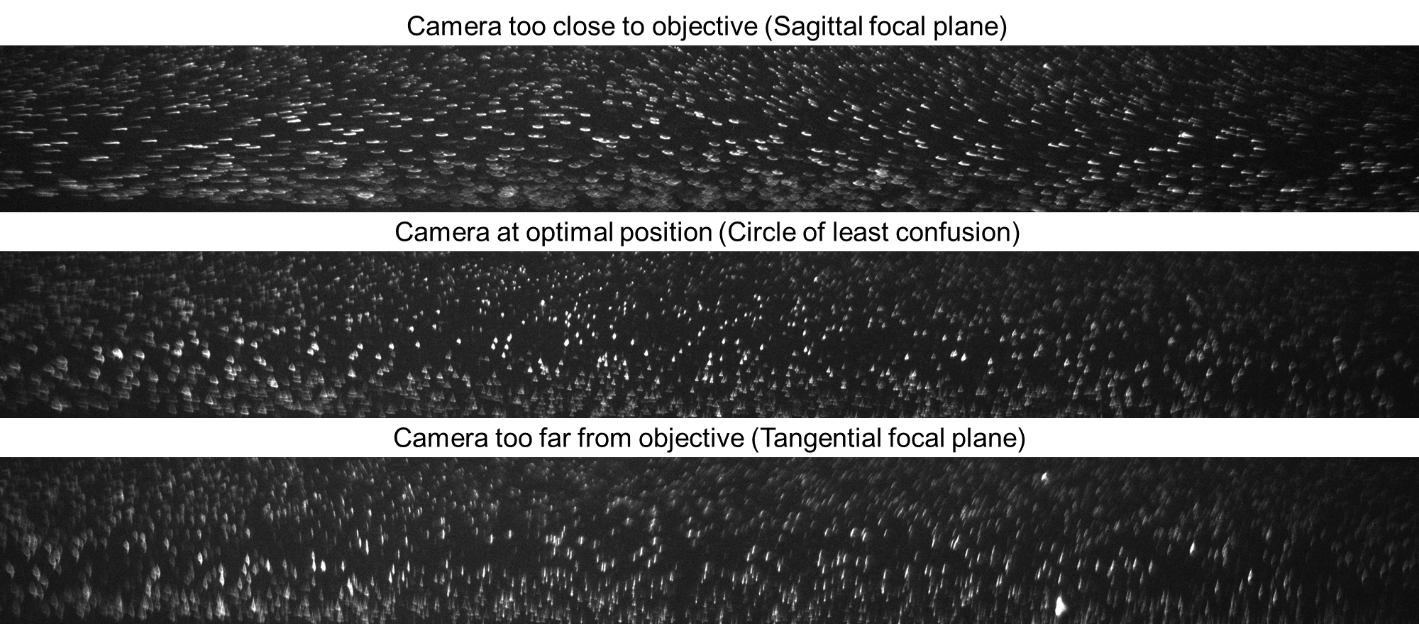

**Supplemental Figure S3. Bead image at different camera positions.** The bead patterns as the camera moves from under- to over-focus positions resemble the airy disk patterns of astigmatism from sagittal to tangential focal planes.

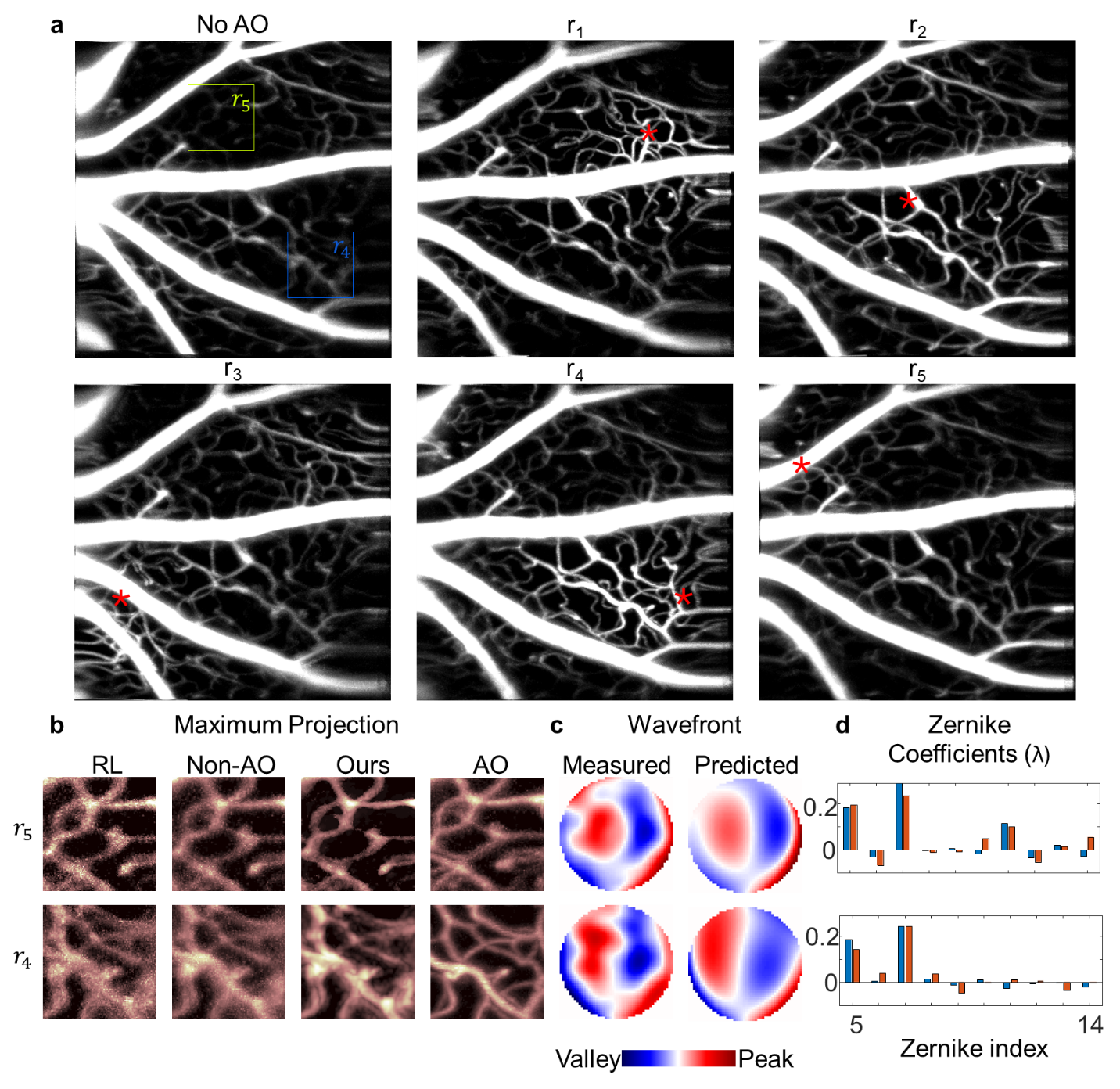

**Supplemental Figure S4.** (a) Mouse retina images with WF sensing applied at different image positions (red asterisks). (b) En face and side view of select ROIs in a (colored squares) from non-AO, RLD, our deconvolved image, and AO-corrected measurements. The experimental SHWFS measurements and aberration predictions for these ROIs are shown as the phase map (c) and Zernike coefficients (d).

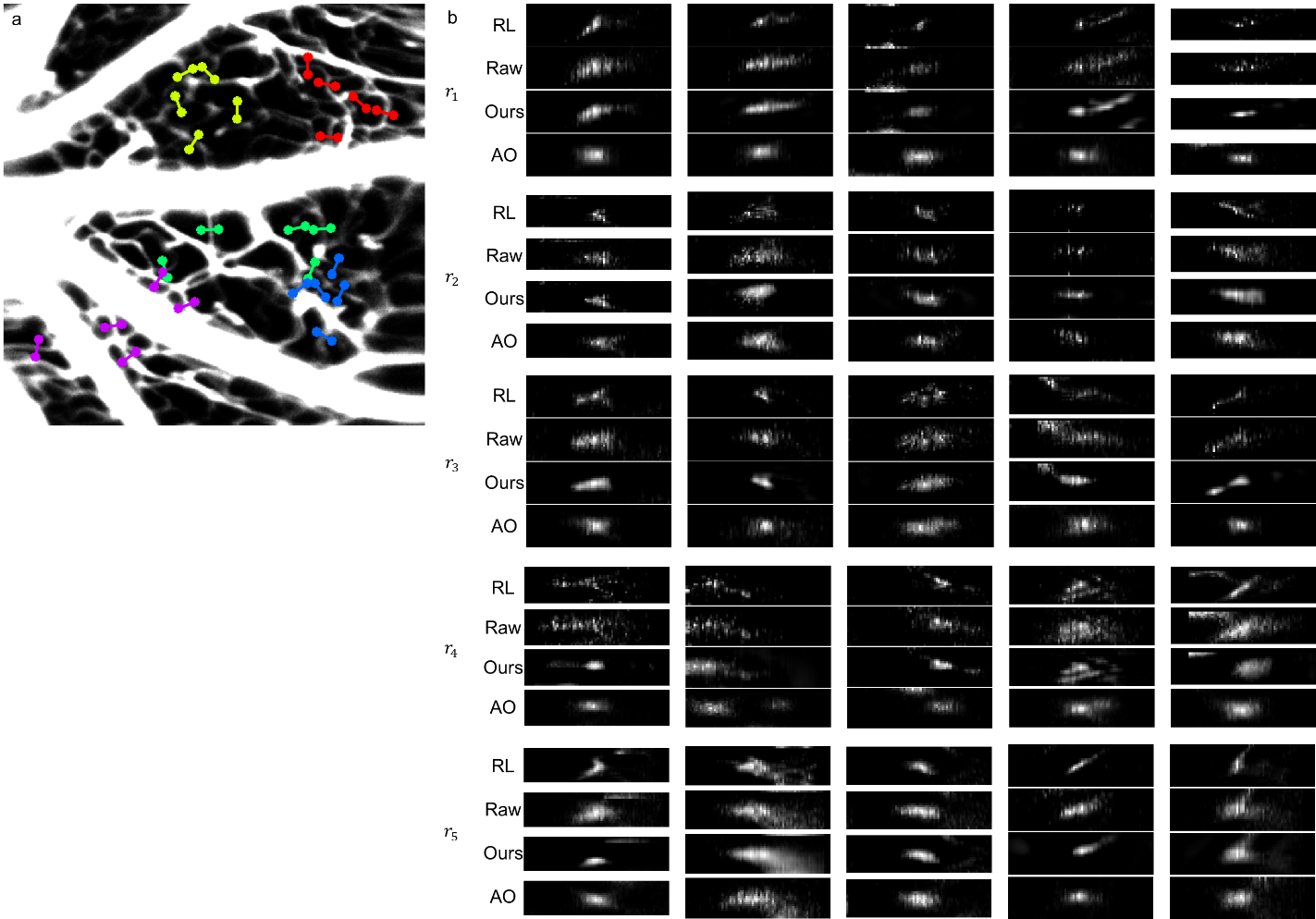

**Supplemental Figure S5. (a) Select capillaries at different AO locations and (b) their cross-sections.**

**Supplemental Table S1. Zernike orders used to represent aberrations.** Each order is a polynomial of the polar pupil coordinates.

| Noll index | Polynomial formula | Classical name |
| --- | --- | --- |
| 5 | $\sqrt{6}\rho^{2}\sin2\phi$ | Oblique astigmatism |
| 6 | $\sqrt{6}\rho^{2}\cos2\phi$ | Vertical astigmatism |
| 7 | $\sqrt{8}\left( 3\rho^{2}-2\rho\right)sin\phi$ | Horizontal coma |
| 8 | $\sqrt{8}\left( 3\rho^{2}-2\rho\right)cos\phi$ | Vertical coma |
| 9 | $\sqrt{8}\rho^{3}\sin3\phi$ | Vertical trefoil |
| 10 | $\sqrt{8}\rho^{3}\cos3\phi$ | Oblique trefoil |
| 11 | $\sqrt{5}\left( 6\rho^{4}-6\rho^{2}+1 \right)$ | Primary spherical |
| 12 | $\sqrt{1}0\left( 4\rho^{4}-3\rho^{2} \right)\cos2\phi$ | Vertical secondary astigmatism |
| 13 | $\sqrt{1}0\left( 4\rho^{4}-3\rho^{2} \right)\sin2\phi$ | Oblique secondary astigmatism |
| 14 | $\sqrt{\left( 10 \right)}\rho^{4}\cos4\phi$ | Vertical quadrafoil |

**Supplemental Table S2: Hyper-parameters for aberration correction.**

| Image | Structure LR | Zernike LR | Epochs | $\lambda_{TV}$ | $\lambda_{L1}$ |
| --- | --- | --- | --- | --- | --- |
| 10x OPM | 5e-2 | 1e-3 | 2000 | 0.1 | 0.1 |
| 20x OPM | 5e-3 | 5e-4 | 2000 | 0.05 | 0.05 |
| AOSLO | 1e-4 | 1e-4 | 1000 | 0.1 | 0.1 |
